# DNA mismatch and damage detection using a FRET-based assay for monitoring the loading of multiple MutS sliding clamps

**DOI:** 10.1101/2021.07.23.453479

**Authors:** Vladislav Kunetsky, Olha Storozhuk, Gwendolyn Brouwer, Charlie Laffeber, Mark S. Dillingham, Joyce Lebbink, Peter Friedhoff

## Abstract

We developed a sensitive, homogeneous fluorescence assay for the detection of DNA mismatches and DNA damage based on the mismatch repair (MMR) protein MutS. The assay is based on Förster resonance energy transfer (FRET) between SYBR Green I (SG), non-covalently bound to DNA, and Alexa Fluor 647 (AF647) conjugated to MutS. In contrast to previous assays using only the mismatch binding activity of MutS, we exploited the ATP-dependent loading of multiple MutS sliding clamps provoked by mismatch/damage to the DNA, which increases the overall sensitivity of the assay. The assay was validated using a well-characterized 3 kb circular DNA containing a single G/T mismatch. We also demonstrate that treatment of long (multiple kb) DNA with various chemical or physical agents including non-denaturing bisulfite conversion of cytosine to uracil, cisplatin modification or ultraviolet light (UVC) results in changes in the DNA that can be detected by the FRET-based MutS biosensor.

## INTRODUCTION

High quality DNA is essential for many applications in molecular biology. However, the manipulation of DNA, beginning with its isolation from a living organism through chemical or enzymatic assembly *in vitro* may lead to damages that are not detected by the standard quality control methodologies currently employed by molecular biologists.

Damage-free samples of very long DNA (>several kb) are particularly difficult to obtain. Quality control of these samples involves at least two steps, i.e. measurement of the quality of DNA and error/damage correction to improve quality (1–3). Recently, next generation sequencing has been used to assess the performance of several error correction enzymes that are currently used to increase the quality of synthetic DNA (2). Our goal was to develop a fast and sensitive tool to assess the quality of long DNA molecules.

We selected the DNA mismatch repair protein MutS from *Escherichia coli*, which not only recognizes mismatches in DNA but can also detect a variety of other DNA damages including base pairs containing O6-methylguanine, 8-oxoguanine, carcinogen adducts, UV photo products, and cisplatin adducts (4). MutS has been used previously as a sensor for DNA mismatches (5–18), including a fluorescence-based method (19), and employed as an error removal enzyme to increase the quality of synthetic DNA (2). However, most assays only exploit MutS binding for the detection of mismatched or damaged DNA. Moreover, when using linear DNA, a further limitation is caused by the DNA end-binding ability of MutS, especially when using short DNA fragments (20). An exception is the MutS, MutL, MutH mismatch detection system, which relies on the mismatch-provoked cleavage of DNA (21–23).

To develop a more sensitive and efficient fluorescence-based assay, we took advantage of two factors not yet exploited by others to increase the overall sensitivity of MutS as a mismatch/damage sensor. First, an ATP-induced change in the MutS-DNA binding conformation: after mismatch recognition ATP switches MutS into a long-lived clamp state placing the DNA in a new binding pocket (24). Second, ATP-induced loading of multiple MutS sliding clamps: the MutS sliding clamp leaves the mismatch allowing the loading of additional MutS at the site of the mismatch/damage (25).

To monitor the conformational change in MutS and the loading of multiple MutS we developed a novel assay that monitors FRET between a donor dye non-covalently attached to DNA and an acceptor dye covalently attached to MutS (Figure 1A). Specifically, we use SYBR Green I (SG), which preferentially stains double-stranded DNA as a donor dye (26) and Alexa Fluor^®^ (AF647) as an acceptor. SG is widely used for real-time PCR and other assays in which the presence of double-stranded DNA is monitored(27). For labelling MutS with AF647 we selected a position in MutS that is far away (> 5 nm) from the DNA in the scanning and mismatch bound states but becomes close to DNA (~3 nm) in the ATP-switched state (Figure 1B). Upon binding of MutS to DNA, a low FRET can occur between DNA-bound SG and single MutS-coupled AF647 and the FRET intensity correlates with the amount of mismatch-bound MutS. After mismatch recognition, ATP triggers the transition of MutS from the mismatch recognition into the sliding clamp state. This pushes the DNA down and closer to the core domain (Figure 1B) resulting in a decrease of the distance between DNA-bound SG and AF647 attached to the core domain (from about 5 nm to 3 nm) and increases the FRET efficiency (24). Most importantly, the MutS sliding clamp leaves the mismatch, which allows loading of multiple MutS (25,28). Consequently, the final FRET signal is amplified compared to that observed from a single mismatch bound MutS, which makes detection of mismatches using this method more sensitive.

**Figure 1.**
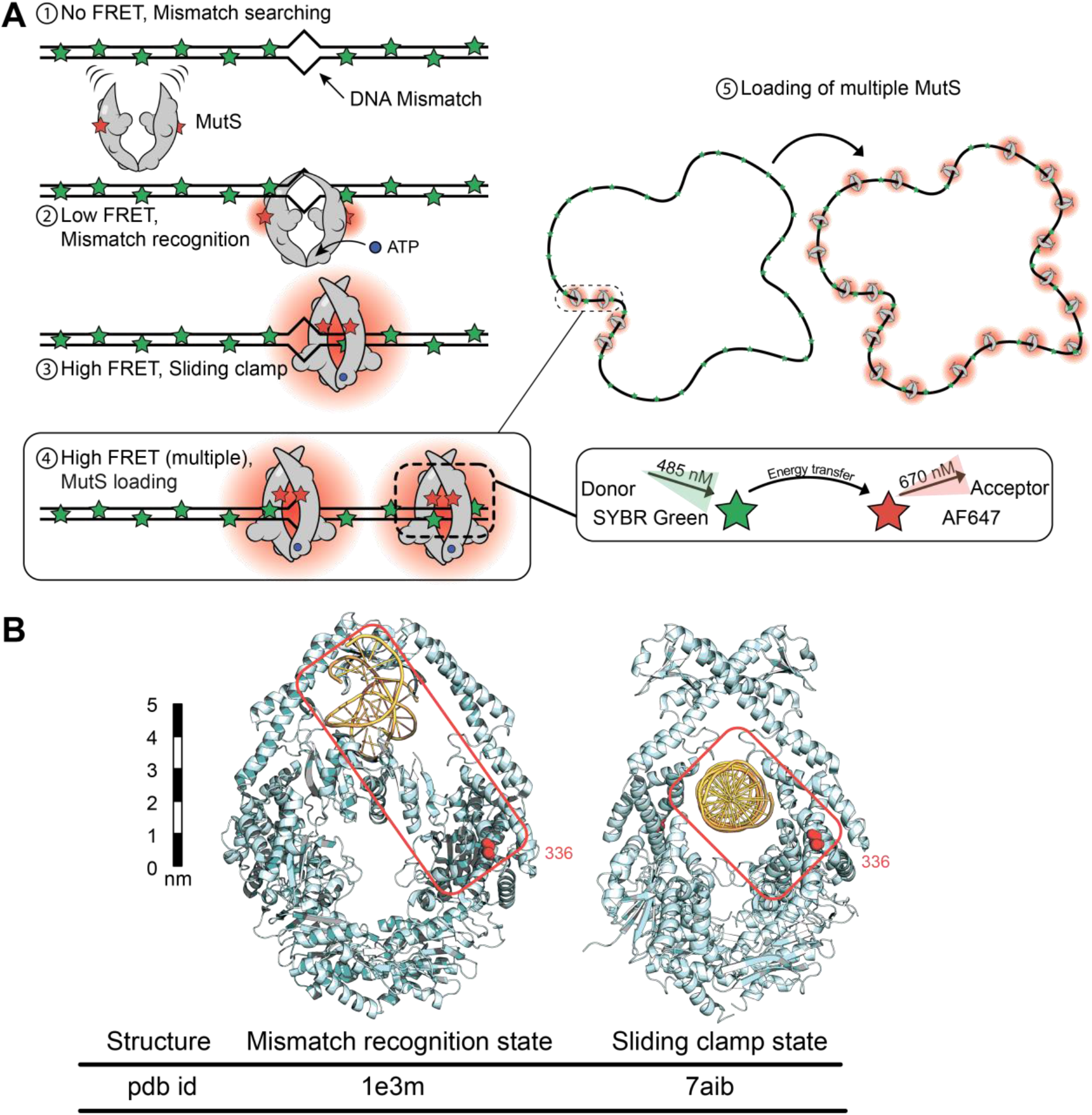
Schematic representation of the FRET-based MutS biosensor assay. **(A)** MutS scans DNA and binds to mismatches in a low FRET mismatch recognition state. ATP binding triggers a conformational change leading to the high FRET MutS sliding clamp state. Sequential binding of additional MutS to the mismatch results in loading of multiple MutS sliding clamps onto the mismatch-containing DNA, which significantly increases the FRET signal. **(B)** MutS structures with amino acid at position 336 shown as red spheres; pdb-codes: 1e3m and 7aib and approximate distances between position 336 in MutS and DNA. In the sliding clamp state, the fluorophore attached to MutS comes closer to the DNA making FRET between DNA-bound SYBR green I (SG) and dye covalently attached to MutS more efficient.

## MATERIAL AND METHODS

### Protein purification

MutS, MutL and UvrD proteins were purified as described previously (24,29). His-tagged MutS dimer variant (MutS336C/D835R) with a single-cysteine at position 336 was created from the dimeric cysteine-free MutS variant (24,30,31) using the NEBuilder^®^ HiFi Assembly Cloning Kit (NEB) (see Supplementary Data for details) and purified as described before (24). Protein concentrations were determined spectrophometrically using theoretical extinction coefficients (ε^280 nm^ =73 605 M^-1^ cm^-1^ for MutS, ε^280 nm^ = 54 270 M^-1^ cm^-1^ for MutL, ε^280 nm^ = 105770 M^-1^ cm^-1^ for UvrD).

His-tagged UvrD was overexpressed in BL21 Star™(DE3) pLysS cells in the vector pET15b and purified by Ni-IDA chromatography. The N-terminal His6-tag was cleaved by TEV protease in dialysis buffer (Tris-HCl 20 mM pH 8.3, NaCl 300 mM, EDTA 1 mM, glycerol 20% v/v, DTT 2 mM) for 14 h at 4°C. Cleaved protein was passed through Ni-TED (Macherey-Nagel) to remove uncleaved protein, His-tag and TEV-protease followed by Heparin, HiTrap Q and desalting columns (Zeba Spin Desalting column 40K). UvrD was stored in storage buffer (25 mM HEPES pH 8.0, 400 mM KCl, and 20% glycerol) and snap-frozen in liquid nitrogen and stored at −70°C.

### Protein labelling

To obtain MutS labelled with Alexa Fluor 647 (AF647-MutS), MutS336C/D835R was diluted to 40 μM in 150 μl of HPLC buffer (10 mM HEPES/KOH (pH 8.0), 200 mM KCl and 1 mM EDTA) and labelled with a 5-fold molar excess of Alexa Fluor 647 maleimide (Invitrogen, Thermo Fisher scientific, Waltham, MA) for 2 h on ice in the dark according to the manufacturer’s instructions. Excess dye was removed using Zeba Spin Desalting columns (Thermo Fisher scientific, Waltham, MA) and the degree of labelling (DOL) was determined from the absorbance spectra (NanoDrop 1000, Thermo Fisher scientific) (DOL 90.3 ± 3.3 %, n=3) as described previously(24). The solution was aliquoted, flash-frozen in liquid nitrogen and stored at −80 °C.

### DNA substrates

The 3196 bp circular G/T mismatch DNA substrate (without fluorescent label) was created by primer extension and ligation as described earlier using the circular single-stranded DNA derived from phagemid GATC1 as a template (32). Plasmid DNAs (phagemid GATC1, 13026 bp (pHis_Tho2) and 15898 bp (pB_Tho2) substrates) from *E. coli* DH5α were purified using the NucleoSpin Plasmid, Mini kit for plasmid DNA (Macherey-Nagel). To obtain linear DNA of the indicated length, plasmid DNA was incubated with the ScaI-HF (New England Biolabs) for 3196 bp Phagemid GATC1 or EcoRI (New England Biolabs) for 13026 bp (pHis_Tho2) and 15898 bp (pB_Tho2), respectively, for 30 min at 37°C followed by heat inactivation for 20 min at 65°C. For 48502 bp linear DNA, λ-DNA (SD0011, Thermo Fisher scientific) was used.

### Non-denaturing bisulfite treatment

For deamination of cytosine bases in DNA under non-denaturing conditions, plasmid DNA was treated using the EpiMark^®^ Bisulfite Conversion Kit (E3318S, New England Biolabs). The protocol was modified to allow non-denaturing deamination. Plasmid DNA (2 μg) was diluted in 100 μl of bisulfite mix and the cycling protocol (95 °C and 60 °C - total incubation time 205 min) was replaced by incubation for 30 min at 50°C (or the indicated temperature) followed by a slowly decreasing temperature after treatment (1°C/min) to room temperature. Excess of sulphonation reagent was removed by using EpiMark™ spin columns according to manufacturers’ instructions. The volume of the eluted DNA solution was adjusted to 150 μl in 1 % desulphonation solution from the kit and incubated for 15 minutes at room temperature. Excess reagent was removed using an EpiMark™ spin column. The eluted, deaminated DNA solution was stored at −20°C.

### UV-irradiation

DNA (plasmid DNA or λ-DNA) was placed at 40 ng/μl in 50 μl in water on parafilm and exposed to UV irradiation at 254 or 365 nm with a controlled UV irradiation system (Bio-Link 254 and Bio-Link 365, Vilber Lourmat Deutschland GmbH, Germany) with 0.8 J/cm^2^. UV-irradiated DNA was stored at 4°C.

### Cisplatin modification

Cis-diammineplatin(II)-dichloride was obtained from Sigma-Aldrich. Plasmid DNA was incubated in 154 mM NaCl water solution in the absence or presence of cisplatin (1.4 μM – 2.8 mM) for 24 h at room temperature in the dark, starting at 2.8 mM with two-fold dilution down to 1.4 μM. Excess cisplatin was removed by using Illustra™ NAP-5 columns (Sigma-Aldrich) for desalting and buffer exchange. Modified DNA was stored at 4°C.

### Nicking of deaminated plasmid

To generate circular DNA with a single nick phagemid DNA (~2,6 μg) with or without non-denaturing bisulfite treatment was treated with 20 Units of Nt.BstNBI (New England Biolabs) for 1 hour at 37°C in 50 μl of 1X NEBuffer 3.1 followed by deactivation at 70°C for twenty minutes.

### Agarose gel electrophoresis

DNA samples (200 ng) were analysed on 0.8% agarose gels in TAE buffer (Tris 40 mM, sodium acetate 20 mM, EDTA 1 mM, pH 8.0) (30 V for 3 hours) stained with HDGreen^®^ Plus Safe DNA Dye and UV fluorescence (Intas Science Imaging Instruments GmbH, Germany).

### MutS sliding clamp formation on DNA monitored by FRET

DNA substrates were diluted to 0.38 ng/μl (575 nM bp; in case of 3.2 kb circular G/T 0.18 nM G/T mismatches) in 200 μl of FB150 buffer (25 mM HEPES-KOH pH 7.5, 5 mM MgCl2, 150 mM KCl, and 0.05 % (v/v) Tween 20) at room temperature in 96-well plates with 250 nM of SYBR Green I (SG) (S7563, Thermo Fisher Scientific) (dye/base pair ratio 0.43) (Invitrogen, Thermo Fisher scientific, Waltham, MA). AF647-MutS (50 nM monomer) was added to the DNA substrates and incubated at room temperature for 150 seconds. ATP (1 mM) was added and kinetics of the fluorescence signals were recorded with a fluorescence microplate reader (TECAN infinite F200, Tecan Group Ltd, Switzerland). Fluorescence intensities were measured in three channels (donor, acceptor, FRET) with the following filter combinations: (donor ex. 450 nm (width 20 nm) em. 535 nm (width 25 nm), acceptor ex. 620 nm (width 10 nm) em. 670 nm (width 25 nm), FRET ex. 485 nm (width 20 mm) em. 670 nm (width 25 nm) filter. Signals were corrected for background from buffer and spectral bleed-through to obtain the corrected signal intensities. Correction factors for spectral bleed-through were obtained from measurements with only one of the labelled components present (see Supplementary Data for details) In particular, the corrected FRET signal FRET^c^ (corrected for background and spectral bleed-through) was calculated using equation (7) (see Supplementary Data) essentially as described before (33).

### Fluorescent SSB binding assay

Single cysteine mutant SSB from *Plasmodium falciparum* was expressed, purified and labelled with N-[2-(iodoacetamido)ethyl]-7-diethylaminocoumarin-3-carboxamide (IDCC) to give DCC-SSB as described earlier(34,35). For measurement of UvrD helicase activity, experiments were performed by preincubating 100 nM of MutS, 100 nM of MutL and 100 nM of UvrD with pre-nicked DNA (0.38 ng/μl) in 200 μl of FB150 buffer (HEPES 25 mM, MgCl2 5 mM, KCl 150 mM, pH 7.5) and 30 nM of fluorescence SSB (DCC-SSB) for 5 min. The reaction was started by rapidly adding 2 μl of 100 mM ATP. Fluorescence intensities were measured with excitation (430 ± 20 nm) and emission (485 nm ± 20 nm) band pass filters.

## RESULTS AND DISCUSSION

### FRET-based MutS biosensor assay using G/T mismatched DNA

We first established the FRET assay using a well-characterized 3.2 kb circular DNA with or without a single G/T mismatch that has been used before to study the initial steps of mismatch repair (32,36) (Figure 2A). The use of a circular rather than linear DNA prevents potential DNA-end binding and dissociation of the MutS sliding clamps. The specificity of the system was verified by using a phagemid DNA of identical sequence but without a mismatch (32,36). In the absence of a mismatch almost no signal (corrected FRET signal; FRET^c^ (37)(see Supplementary Data for details), 1300 ± 200 a. u. Figure 2B) was detectable in the FRET channel after correction for spectral crosstalk (see Material and methods and Supplementary Data). This was independent of the absence or presence of ATP (Figure 2B column 1 +2). For the G/T mismatch DNA in the absence of ATP, the FRET^c^ signal was small (2400 ± 400 a. u.) but significantly higher (1.8 fold) compared to that with DNA without a mismatch (Figure 2B column 1+3). Upon addition of ATP, a substantial (> 12-fold) increase of the FRET^c^ was observed for G/T DNA (28000 ± 3000 a.u.) (Figure 2B column 4) confirming our assumption that loading of multiple MutS sliding clamps would lead to a large signal amplification. In comparison to the plasmid control with ATP this increase is 24-fold (Figure 2B column 4 vs. column 2).

**Figure 2.**
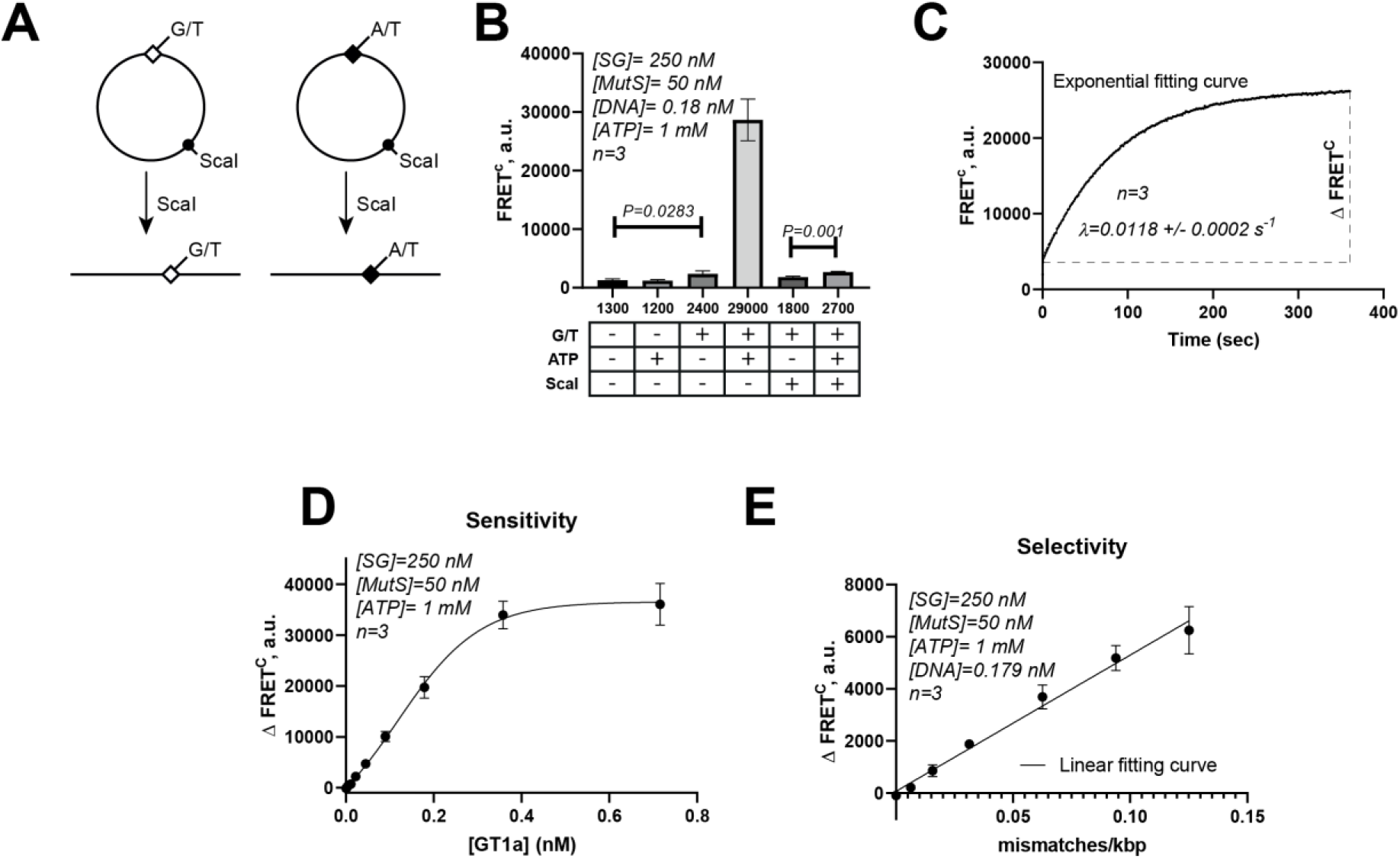
Mismatch detection by the FRET using MutS **(A)** Schematic representation of the DNA **(B)** FRET^c^ between AF647-MutS and SG-stained mismatch or phagemid DNA substrates. **(C)** Kinetics of loading multiple MutS on the circular G/T mismatch DNA. **(D)** Analysis of sensitivity: Concentration dependence of circular G/T mismatch DNA on ΔFRET^c^ (FRET^c^_ATP_ - FRET^c^_no ATP_). **(E)** Analysis of selectivity: Decreasing fraction (expressed as G/T / kb) of circular G/T mismatch DNA in the presence of constant amount total DNA ([G/T mismatch DNA] + [phagemid] = 0.179 nM and SG (250 nM) keeping a constant dye/ base pair ratio (dbpr) of 0.44. Note, the G/T mismatch DNA has a G/T per kb of 0.31.

Under the conditions used, half-maximum signals were achieved for SG at 140 nM, AF647-MutS at ≈ 30 nM and ATP at ≈ 7 μM (Supplementary Figure S1). Since MutS sliding clamps can diffuse at least 1 kb along the DNA backbone and dissociate from DNA via its ends(25), we asked whether linearization of the 3 kb circular G/T mismatch DNA would reduce the number of MutS sliding clamps on DNA and hence the observed signal for FRET^c^ would decrease. Cleavage of the circular G/T by ScaI results in a linear form with a 1.4 kb arm on the 5’-side and a 1.7 kb arm on the 3’-side of the G/T mismatch. In the absence of ATP, the FRET^c^ was only slightly reduced (Figure 2B column 4, non-significant, *p* = 0.15), indicating that linearization did not affect the binding of MutS to the G/T mismatch. However, addition of ATP resulted in only a small (1.5-fold) increase of FRET^c^ (*p* = 0.01), much smaller (by > 85 %) compared to the FRET^c^ increase observed with the circular form of the G/T-mismatch DNA. This suggests that on the linear G/T DNA a strongly reduced number of sliding clamps are present (Figure 2B column 5).

Next, we investigated the kinetics of the ATP-induced loading of multiple MutS sliding clamps on the circular G/T mismatch DNA. Stopped-flow kinetics revealed biphasic kinetics with a rapid burst (~1 s) followed by a slower exponential phase (Supplementary Figure S2). Upon addition of ATP the FRET^c^ increased over time and a single exponential function could be used to fit the data of the second phase with an amplitude ΔFRET^c^ of about 23000 (a.u.) and a decay rate λ of about 0.01 s^-1^ (lifetime τ ≈ 100 s) (Figure 2C and Supplementary Figure S3). This is in the same order of magnitude as the reported lifetimes of a MutS sliding clamp on DNA with blocked-ends using surface plasmon resonance (≈ 70 s)(38) further suggesting that our assay monitors mismatch-induced loading of multiple MutS sliding clamps.

To determine the sensitivity of the assay, we quantified the ATP-induced ΔFRET^c^ as a function of the concentrations of circular G/T mismatch DNA (Figure 2D). A linear range was obtained from 0.01 nM to 0.2 nM [GT1a]. We then changed the ratio of mismatches to total DNA at a fixed concentration of total DNA (570 nM bp) by mixing GT1a with phagemid DNA. The circular G/T mismatch DNA has a density of 0.33 mismatch per kb. Under these conditions, the limit of detection (LoD was approximately 1 mismatch per 240 kb (or 0.004 mismatches/kb) and the limit of quantitation (LoQ) was 1 mismatch per 50 kb (or 0.02 mismatches/kb) (Figure 2E and Supplementary Figure S4).

Although loading of multiple MutS sliding clamps was enhancing the sensitivity of mismatch detection, a limitation seemed to be the requirement for circular DNA. It is well-established that during MMR a ternary complex between DNA, MutS and MutL is formed in a mismatch and ATP-dependent manner(25,39) and that the MutS-MutL-complex slides more slowly compared to MutS alone(24,25,39,40). Using electrophoretic mobility shift assays, at a concentration of 100 nM MutS and 100 nM MutL, a significant but incomplete reduction of the fraction of complexes bound to 2.9 kb of G/T mismatch DNA after linearization was observed(25). Consequently, we asked whether and how MutL influences the signal for FRET^c^ from MutS sliding clamps on circular and linear 3 kb G/T mismatch DNA. While on circular DNA, MutL had little effect on the signal for FRET^c^ (Figure 3A), on the G/T mismatch DNA linearized with ScaI we observed a strong dependence on FRET^c^-signal on the concentration of MutL (Figure 3B). A hyperbolic function was fitted to the data with an apparent *K*1/2 of about 50 nM. This value is similar to that reported before for *Thermus aquaticus* MutL trapping MutS on DNA(41).

**Figure 3.**
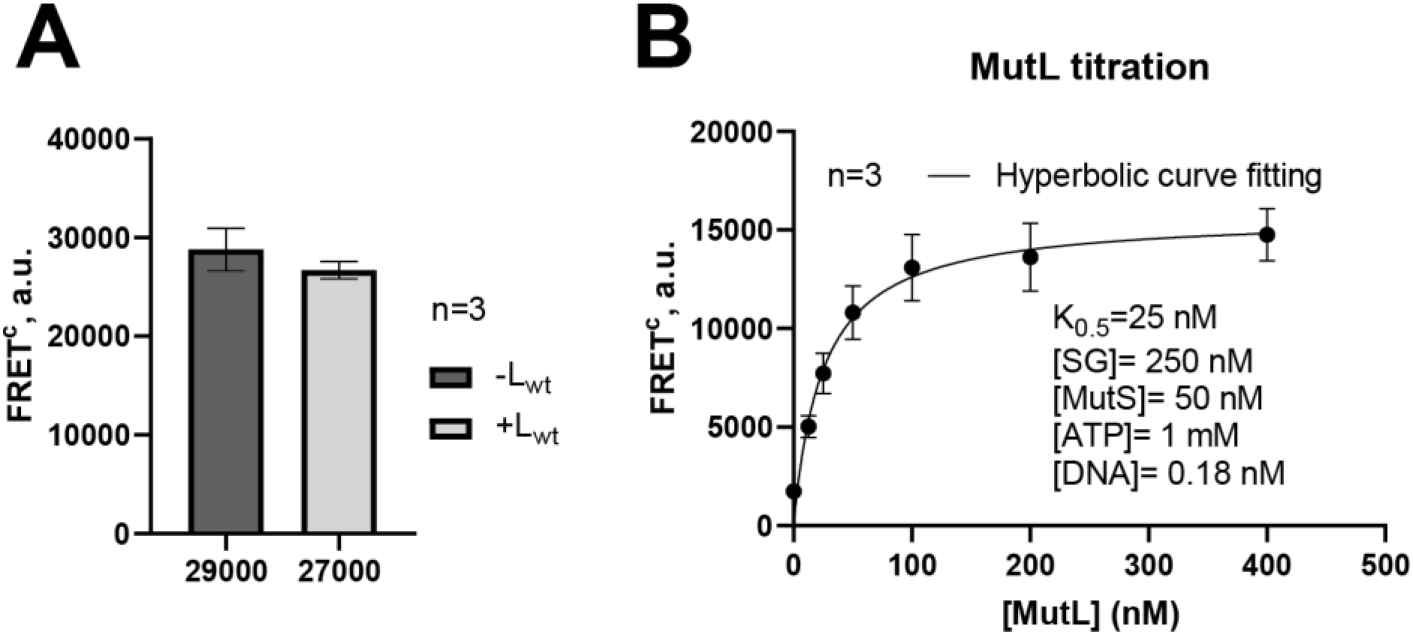
Mismatch detection on by FRET using MutS and MutL **(A)** Influence of MutL on the observed FRET^c^ with AF647-MutS and circular G/T DNA **(B)** Concentration dependence of MutL on the observed FRET^c^ between MutS-AF657 and linear G/T DNA.

### Detection of DNA deamination by MutS FRET Biosensor

Taken together, our data suggest that labelled MutS has the potential to act as a biosensor for certain DNA damages. To demonstrate the ability of our assay to detect DNA damage, we first exploited the ability of MutS to bind to G/U mismatches that can arise from spontaneous or enzymatic deamination or via misincorporation of dUMP into DNA(42). We used the phagemid GATC1, which contains a cruciform forming sequence that shows enhanced reactivity towards bisulphite-induced deamination of cytosine under non-denaturing conditions (43). Since spontaneous deamination in double-stranded DNA is rather slow, we changed a protocol for cytosine deamination using the NEB EpiMark^®^ Bisulfite Conversion Kit to milder conditions (see Materials and Methods and Supplementary Figure S5) to convert circular plasmid DNA into G/U containing DNA.

To verify that the bisulfite conversion (BC) did indeed result in the deamination of cytosine to uracil, Uracil-DNA glycosylase (UDG) was used to introduce apyrimidinic (AP) sites by excision of the uracil base, and then endonuclease Endo IV to convert the AP site to a single-strand. This transforms the supercoiled, covalently closed phagemid DNA into open circular DNA, which can be detected by agarose gel electrophoresis. In the absence of bisulfite conversion, neither treatment with Endo IV nor Endo IV/UDG resulted in the formation of the open circular form (Figure 4A, lane 2+3). Bisulfite conversion at 50 °C alone, or followed by Endo IV treatment, also did not result in formation of the open circular form (Figure 4A lane 4+5). Only BC, Endo IV and UDG treatment resulted in the formation of an open circular form of the plasmid DNA (Figure 4A lane 6) suggesting that uracil was indeed present in the bisulfite-treated DNA.

**Figure 4.**
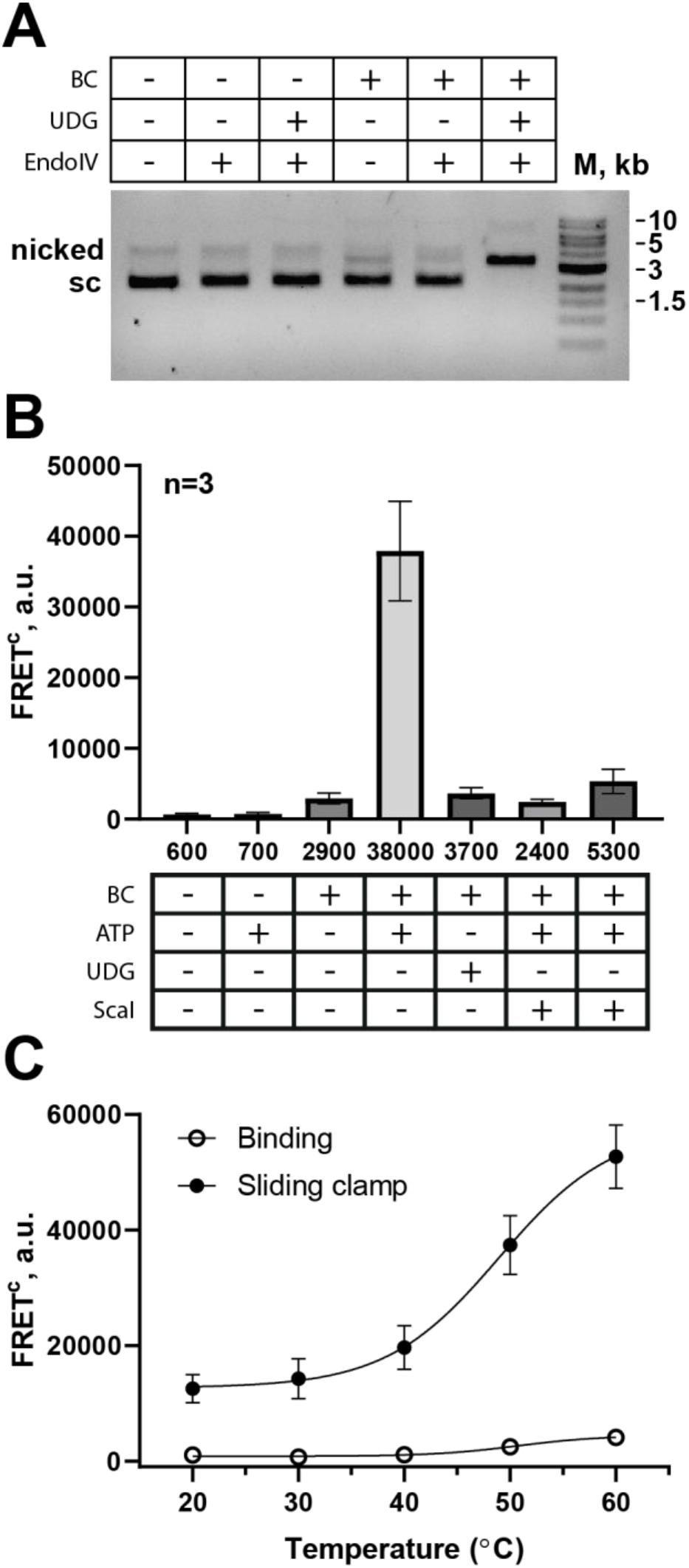
Bisulfite conversion of phagemid DNA can be monitored by the MutS biosensor (**A**) Bisulfite treatment of phagmid DNA under non-denaturing conditions at 50 °C leads to conversion of cytosine to uracil detected by combined UDG/EndoIV treatment and separation of nicked from supercoiled DNA on agarose gel. (**B**) Bisulfite conversion under non-denaturing conditions leads to loading of multiple MutS sliding clamps as detected by FRET^c^. (**C**) Dependence of the FRET^c^-signal on the temperature during bisulfite conversion under non-denaturing conditions.

Next, we tested whether treatment of the phagemid DNA resulted in changes that can be detected by our MutS biosensor. Qualitatively, the results are comparable to those obtained with the G/T mismatch DNA (Figure 2B and 4B). However, the intensity of the signal for FRET^C^ both in the absence (2900 ± 120 a.u.) and presence of ATP (38000 ± 5700 a.u.) was higher after bisulfite conversion DNA (at 50°C) compared to the circular DNA with a single G/T mismatch suggesting the presence of multiple G/U mismatches on at least a fraction of the DNA molecules.

It has been shown for the eukaryotic homologs of MutS that binding to a DNA with an AP site is strongly reduced compared to G/U mismatched DNA(44). Consequently, we asked how uracil excision of the bisulfite-treated DNA by UDG would influence the FRET^c^-signal. Indeed, after UDG-treatment the FRET signal was strongly (10-fold) reduced compared to the untreated DNA suggesting that an AP-site is, at best, a poor substrate for MutS sliding clamp formation compared with G/U containing DNA (Figure 4B).

As with the G/T mismatch DNA, the FRET signal was significantly lower (14 %) for DNA linearized compared to the circular form of the DNA (Figure 2B and Figure 3A). Next we investigated the influence of temperature (between 20 °C and 60 °C) on deamination during bisulfite-treatment and measured the FRET^c^ to test for MutS binding and sliding clamp formation on the circular phagemid DNA (Figure 4B). At temperatures as low as 20 °C the phagemid DNA was readily converted into a substrate capable of inducing MutS sliding clamp formation, although to a lesser extent than G/T mismatch DNA. When the temperature was raised further, the extent of MutS sliding clamp formation was increased. Analysis of the bisulfite-treated DNA by agarose gel electrophoresis revealed no obvious changes compared to the untreated DNA (Figure 4C).

Next, we tested whether the bisulfite-treated phagemid DNA is a substrate for mismatch repair unwinding reactions (Figure 5A). To monitor unwinding we took advantage of a fluorescently labeled single-stranded binding protein (DCC-SSB) that had been developed before as as a sensor for DNA unwinding (34,35). Binding of DCC-SSB to single-stranded DNA results in significantly increased fluorescence (Figure 5A). Using a prenicked circular DNA substrate (Figure 5A) treated with bisulfite we were indeed able to observe the kinetics of DNA unwinding (Figure 5B). The signal was dependent on bisulfite-treatment, a nick, MutS, MutL, UvrD and ATP (Figure 5C).

**Figure 5.**
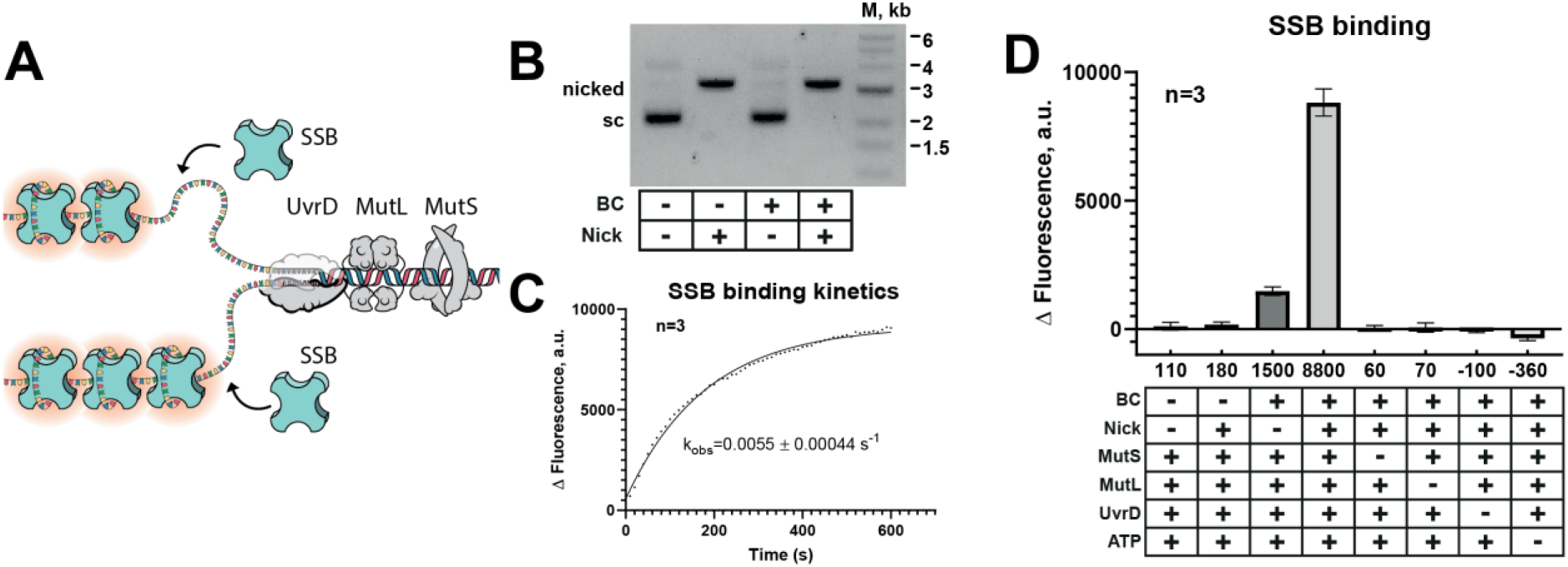
Bisulfite conversion of phagemid DNA leads to activation of MMR **(A)** Schematic representation of Fluorescence-based DCC-SSB binding assay for monitoring UvrD unwinding activity in MM dependent manner. **(B)** Nicking of treated and untreated DNA by Nt.BstNBI. **(C)** Kinetics of DCC-SSBs binding to the ssDNA **(D)** Fluorescence upon binding DCC-SSB to the DNA substrates treated as indicated.

### Detection of damages induced by cisplatin modification

Cisplatin cytotoxicity results from the formation of lesions that block DNA polymerase and cause replication arrest. The damages induced by cisplatin are commonly repaired by nucleotide excision repair (NER)(45). It was shown that *E. coli dam*- strains are more susceptible for the cytotoxic action of cisplatin and N-methyl-N’-nitro-N-nitrosoguanidine (MNNG) than wild type(46–50). Replication arrest can cause damages such as daughter-strand gaps and double-strand breaks. *In vitro* experiments have shown that MutS probably recognizes cisplatin-intrastrand crosslinks, but the specificity is much lower compared to mismatches(51). Here we used our novel FRET-assay to test whether the treatment of DNA with cisplatin lead to damages that are recognized by MutS, leading to the loading of multiple MutS sliding clamps. Indeed, the amplitude of the FRET^c^ signal increased with increasing cisplatin concentration during DNA treatment (Figure 6A). At high cisplatin concentration (> 0.1 μM) the FRET^c^ signal was clearly increased even in the absence of ATP indicating large numbers of lesions recognized by MutS binding. A further significant increase was observed upon adding ATP due to sliding clamp formation of MutS. In contrast to the bisulfite-treated DNA, the difference between binding (no ATP) and sliding clamp formation (ATP) was much lower, suggesting that the lesion introduced into the DNA upon cisplatin-treatment might not convert MutS into sliding clamps efficiently. Similar results were observed with λ-DNA (Supplementary Figure S6). As a quality control, the cisplatin-DNA treated DNA was analyzed by agarose-gel electrophoresis (Figure 6B). At concentrations above 0.36 mM cisplatin changed the electrophoretic mobility of the DNA.

**Figure 6.**
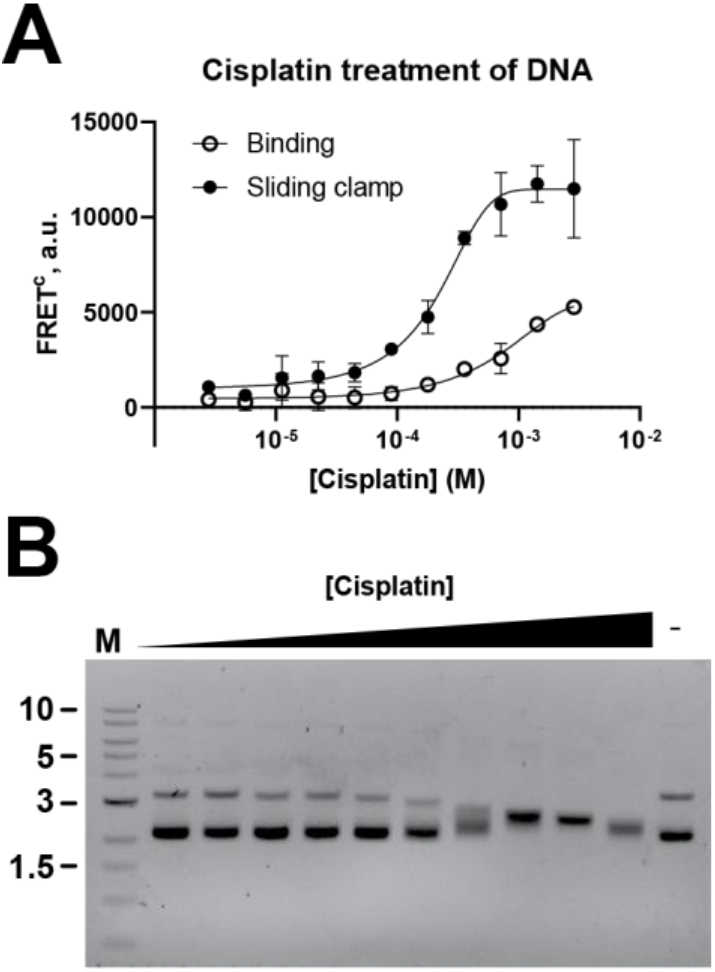
Cisplatin treatment of DNA leads to binding and loading of multiple MutS sliding clamps. (**A**) Dependence of FRETc-signal in the absence (binding) or presence of ATP (sliding clamp) on the concentration of cisplatin during treatment of DNA. (**B**) Analysis of phagemid DNA treated with increasing concentrations of cisplatin by agarose gel electrophoresis.

### Detection of damages induced by UV light

MMR participates in non-canonical recognition of different lesion types including damages induced by UV(52). Moreover, it was shown that human MutSβ is recruited to the sites of UV damages in human (HeLa, XPA, U2OS) cells (53,54). Therefore, we investigated whether our FRET-based assay for detection of multiple MutS sliding clamps could also detect UV-damages in DNA. Indeed, with increasing doses of UV radiation (254 nm), the amplitude of the FRET^c^ signal increased both for MutS binding (no ATP) and loading of multiple MutS sliding clamps (Figure 7A). In contrast to UV radiation at 254 nm, we did not observe any increase in FRET^c^ signal upon treatment with UV 365 nm radiation (Figure 7A). Analysis of the DNA treated with UV radiation of increasing energy by agarose gel electrophoresis showed changes in the electrophoretic mobility even at the lowest energy (0.1 Joule/cm^2^). At the highest energy used (0.8 Joule/cm^2^) virtually all DNA that was originally migrating at the position of supercoiled DNA was shifted to lower electrophoretic mobility. Changes in DNA structure caused by UV damages have also been observed before using atomic force microscopy (55,56).

**Figure 7.**
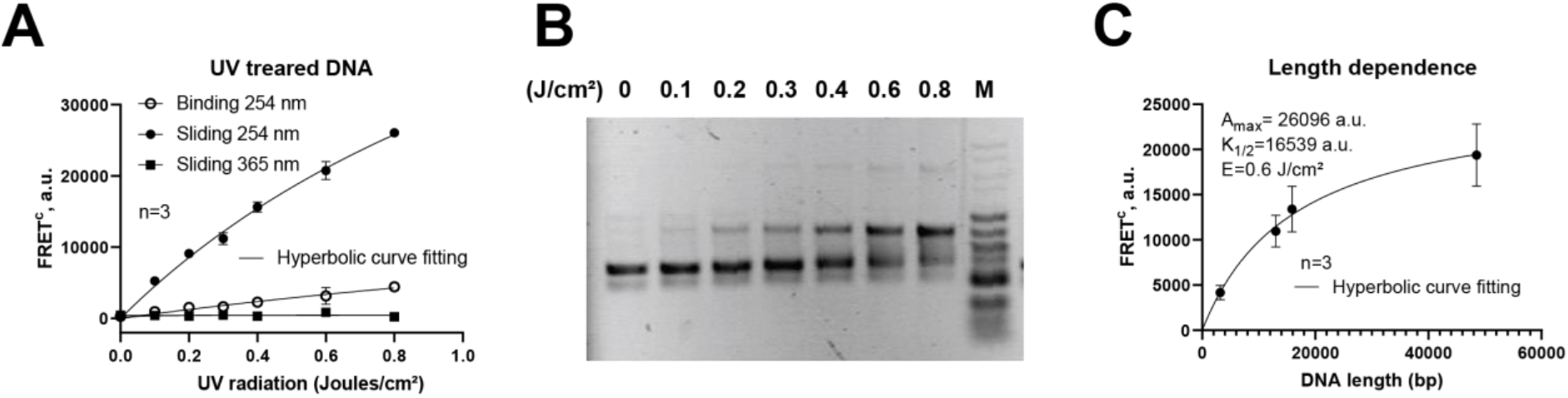
UV treatment of DNA leads to binding and loading of multiple MutS sliding clamp. **(A)** Phagemid DNA was treated with UV light (254 or 365 nm) at the indicated total energy, and loading of multiple AF647-MutS sliding clamps to SG-stained DNA was monitored using FRET. **(B)** Analysis of phagemid DNA irradiated at 254 nm with the indicated energy of UV radiation by agarose gel electrophoresis. **(C)** Length dependence of linear DNA UV-treated (254 nm; 0.6 Joules/cm^2^) substrates for the loading of multiple MutS sliding clamp.

As mentioned above, on linearized 3 kb phagemid DNA containing a single G/T mismatch we did not observe significant loading of multiple MutS sliding clamps (Figure 2B) which was most likely a consequence of rapid dissociation of MutS from the DNA-end via linear diffusion. Using UV-treatment of linear DNA, we reinvestigated this phenomenon using DNA of different length (Figure 7C). First, we were able to observe the loading of multiple MutS sliding clamps onto the linearized 3 kb phagemid DNA treated with UV (0.6 Joule) which was 20 % of that on the circular form but significantly higher (2-fold) than the linearized form of the phagemid DNA with a single G/T mismatch (Figure 2B). Moreover, upon increasing the length of the UV-treated (at constant energy) DNA up to 48 kb (*λ*-DNA) the signal increased. This is in good agreement with the diffusion coefficient of MutS sliding clamps on DNA of 0.0034 μm^2^s^-1^ (39), as it takes about less than 10 s for MutS to travel 3 kb, 300 s for 15 kb and 3000 s for 48 kb (Figure 7C). As the lifetime of MutS sliding clamps is on the order of 100 – 200 s(39), MutS is unlikely to reach the ends of 48 kb DNA within the timescale of the assay.

## CONCLUSION

Here, we present a microliter-plate fluorescence assay which is suitable for quantitative and rapid DNA quality control, in particular to check for mismatches and certain other types of DNA damages in kilobase pair long DNA. In common with several other mismatch detecting approaches used in the past, we took advantage of the mismatch binding protein MutS. The assay is based on FRET between a MutS variant labelled with Alexa Fluor 647 as the acceptor dye and DNA non-covalently stained with the widely used and well-described intercalating dye SYBR Green I (SG), making the method applicable to a wide range of natural or synthetic DNA molecules without the need for labelling. Unlike other methods that rely only on the mismatch binding activity of MutS, we exploited our biochemical and structural understanding of the MutS sliding clamp state in order to increase the sensitivity and specificity of the assay in detecting mismatches in millions of perfectly matched base pairs. The ATP-triggered conformational change and subsequent loading of multiple MutS proteins onto DNA containing mismatches or damages strongly amplifies the signal compared to methods based on binding alone (57). Possible limitations of this signal amplification on shorter DNA were overcome by inclusion of the MutS natural binding partner MutL. Blocking DNA ends e.g. by end binding proteins such as AddA^*H*^B (58) or using roadblocks such as dCas9 could be an alternative (59). We applied this assay for several types of mismatches or damage: (i) GT mismatches which appear during replication, (ii) the effect cytosine deamination, a process which takes place in the cell and can occur during manipulation with DNA for biotechnical purposes, (iii) cisplatin modification or (iv) high energy UV-damages. Although our method does not allow one to detect all kind of damage, it adds to the tools available for damage detection and DNA quality control in *vitro.*

## ACKNOWLEDGEMENTS

We thank Heike Büngen for the excellent technical assistance.

