## Supplementary Information for "DNA mismatch and damage detection using a FRET-based assay for monitoring the loading of multiple MutS sliding clamps"

### SUPPLEMENTARY DATA

#### Cloning of His-tagged single-cysteine MutS-A336C/D835R dimer variant

Single-cysteine MutS (A336C)/D835R was generated by site-directed mutagenesis using HiFi-DNA assembly reaction (NEB). As a template for PCR the plasmid carrying the gene for the cysteine-free (MutS-CF/D835R) was used(1) and the following primer pairs (SePOP purified, Eurogentec): For the linearized vector (5751 bp) MutS_3UTR_Sense (25-mer): gctagcgctatatgcgttgatgcaa and MutS-5UTR_antisense (25-mer): tctagaggggaattgttatccgctc. For fragment 1 (1872 bp) MutS_A336C_sense (27-mer): gatttcaccTGCgggctacagccggta and MutS-3UTR (25-mer): ttgcatcaacgcatatagcgctagc. For fragment 2 (1134 bp) MutS-5UTR (25-mer): gcggataacaattcccctctagaaa and MutS_A336C_antisense (30-mer): tgtagcccGCAggtgaaatcctgcaatgcg. Gene and promotor region were verified by Sanger DNA sequencing.

### Calculation of FRET signals

FRET signal was corrected for spectral bleed-through and spectral crosstalk to obtain the correct signal intensities. Initially, the spectral bleed-through coefficients $S_1$-$S_4$ were obtained from experiments with only a donor or an acceptor by applying next formulas (2,3).

| variable | name | description |
|---|---|---|
| $D_d$, $D_a$, $D_{da}$ | donor channel signal | background-corrected signal in the donor channel for sample that contains donor, acceptor or both, respectively |
| $F_d$, $F_a$, $F_{da}$ | FRET channel signal | background-corrected signal in the donor channel for sample that contains donor, acceptor or both, respectively |
| $A_d$, $A_a$, $A_{da}$ | Acceptor channel signal | background-corrected signal in the donor channel for sample that contains donor, acceptor or both, respectively |
| $S_1$, | bleed-through factor 1 | describes spectral bleed-through from donor into FRET |
| $S_2$ | bleed-through factor 2 | describes spectral bleed-through from acceptor into acceptor channel |
| $S_3$ | bleed-through factor 3 | describes spectral bleed-through from donor into acceptor channel |
| $S_4$ | bleed-through factor 4 | describes spectral bleed-through from acceptor into donor channel |
| $D^c_{da}$ | corrected donor signal | Signal in donor channel, corrected for spectral bleed-through |
| $A^c_{da}$ | corrected acceptor signal | Signal in donor channel, corrected for spectral bleed-through |
| $FRET^c$ | corrected FRET signal | FRET value corrected for spectral bleed-through according to Youvan et al. (4) |
| $N_{FRET}$ | Normalized FRET | normalized FRET measure according to Xia et al (5) |

*Donor into FRET channel:*

$$S_1 = \frac{F_d}{D_d} \quad (1)$$

*Acceptor into FRET channel:*

$$S_2 = \frac{F_a}{A_a} \quad (2)$$

*Donor into acceptor channel:*

$$S_3 = \frac{A_d}{D_d} \quad (3)$$

*Acceptor into donor channel:*

$$S_4 = \frac{D_a}{A_a} \quad (4)$$

The signal values of the donor and acceptor channels containing the mixture in which both components are present has been corrected for spectral bleed-through using next equation

$$D_{da}^{c} = \frac{D_{da} - S_4 \cdot A_{da}}{1 - S_3 \cdot S_4} \quad (5)$$

$$A_{da}^{c} = \frac{A_{da} - S_3 \cdot D_{da}}{1 - S_3 \cdot S_4} \quad (6)$$

In the case of a FRET pair of SG and AF647 fluorophores, the donor fluorescence does not give any signal in acceptor channel and vice versa, meaning that the bleed-through factor 3 and 4 will be close to zero and the corrected donor and acceptor signals will not be changed much.

To obtain actual FRET signal ($FRET^c$), the intensity of the corrected donor and acceptor signals were multiplied by the corresponding spectral bleed-through factors and subtracted from FRET channel signal.

$$FRET^C = F_{da} - D_{da}^C \cdot S_1 - A_{da}^C \cdot S_2 \quad (7)$$

To normalize FRET signal a formula suggested by Xia(6) was used, which normalize FRET signal to the square root of the product of donor and acceptor signal.

$$N_{\mathrm{FRET}} = \frac{FRET^C}{\sqrt{D_{da}^C \cdot A_{da}^C}} \quad (8)$$

**Supplementary table 1**. Filters[1] and bleed-through coefficients.

| ***Donor channel*** | | ***FRET channel*** | | ***Acceptor channel,*** | | $S_1$ | $S_2$ | $S_3$ | $S_4$ |
|---|---|---|---|---|---|---|---|---|---|
| **Ex** | Em | Ex | Em | Ex | Em | | | | |
| **450 (20)** | 535 (25) | 485 (20) | 670 (25) | 620 (10) | 670 (25) | $3.39 \pm 0.07 * 10^{-1}$ | $6.4 \pm 0.11 * 10^{-2}$ | ≈ 0 | ≈ 0 |

[1]center wavelength (bandwidth) in nm

Crosstalks for heated DNA at different temperatures differ from other DNA and depends on the fraction of ssDNA. The emission spectra of DNA stained by SG shifts in the presence of the ssDNA.

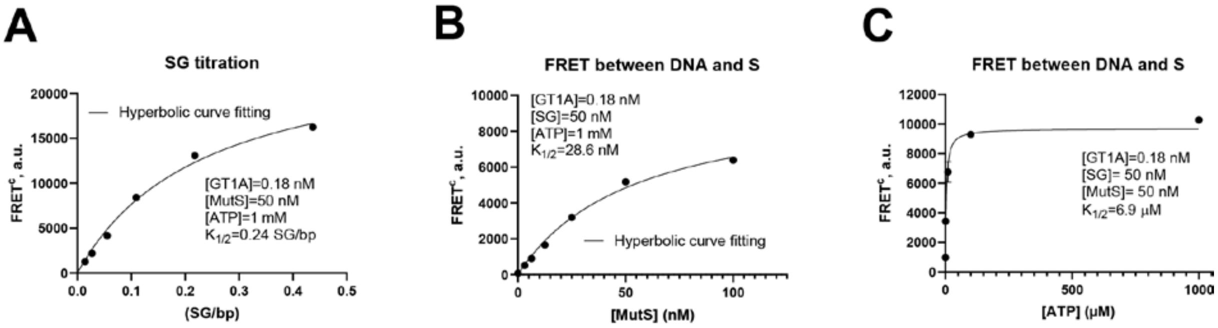


**Supplementary Figure S1.** Concentration dependence **(A)** on SG, **(B)** AF647-MutS or **(C)** ATP on the observed FRET signal intensity on circular mismatched DNA

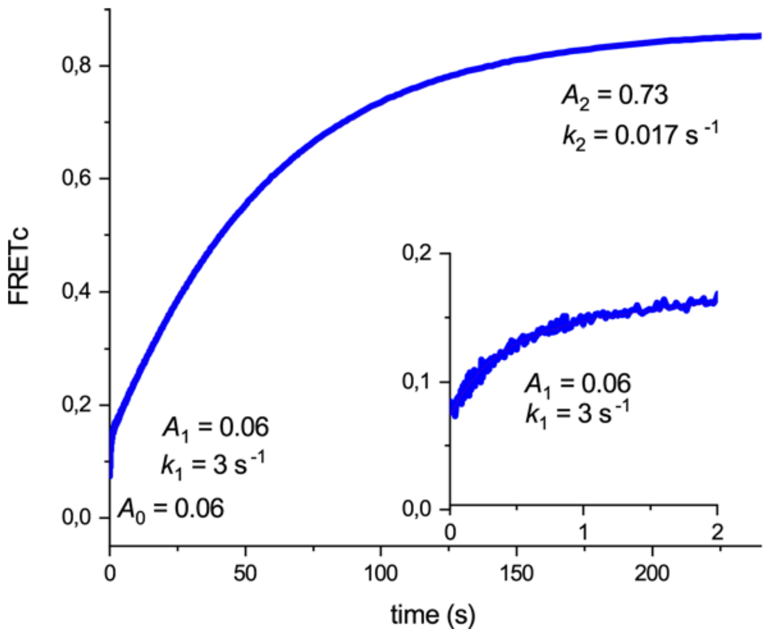


**Supplementary Figure S2.** Stopped-Flow kinetic of ATP-induced loading of multiple MutS sliding clamps. AF647-MutS (100 nM) and 0.36 nM G/T mismatch DNA in FB150T (25 mM HEPES pH 7.5, 5 mM 500 nM SG was mixed with an equal volume of ATP (2 mM) in FB150T. A double exponential function was used to fit the data. The initial values of $FRET^c$ is due to the single mismatch bound MutS. The burst amplitude $A_1$ of 0.06 almost doubles the $FRET^c$ in agreement with the ATP-induced change from a low FRET to a high FRET state (see Figure 1B). The amplitude $A_2$ of the slow phase of 0.73 is best explained with the loading of additional (5-6) MutS sliding clamps in the high FRET state.

Fast kinetics were measured using a stopped-flow device SF-61SX2 (TgK Scientific, Bradford-on-Avon, UK) with an LED light source LSM-470A (Ocean Optics BV) with an in-line filter holder and a long pass filter 475 nm and a short pass filter 500 nm (Edmund Optic® Europe), filter ET525/50 M (Chroma Technology, Olching, Germany) for the donor fluorescence signal and a ET670/50M (Chroma Technology, Olching, Germany) for FRET signal at 25 °C.

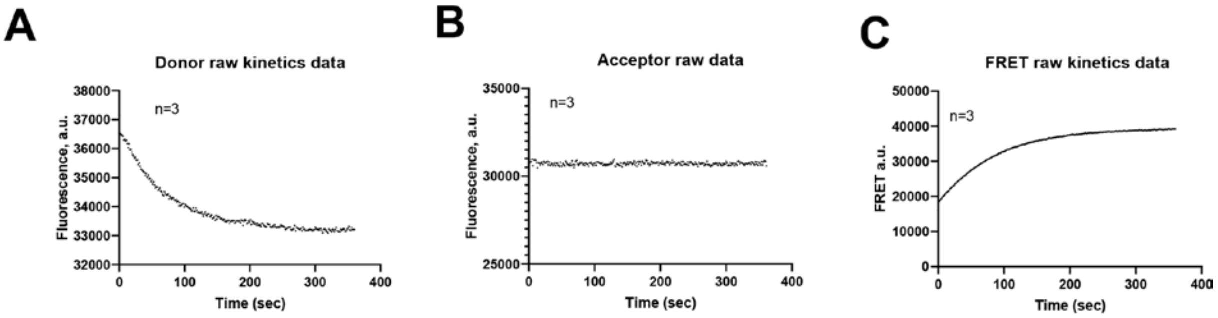


**Supplementary Figure S3. Raw data of kinetics for donor (A) Acceptor (B) and FRET (C) channels**. Raw data of kinetics shown in Figure 2C for donor (A) acceptor (B) and FRET (C) channels. As time increases, the number of proteins that are on the DNA increases, reaching equilibrium after five minutes. (A) As the number of complexes increases, a decrease in donor signal (from 37000 to 33000 a.u.) is observed, as energy is transferred from the donor to the acceptor in the form of FRET. (B) When the acceptor is directly excited, no change in the signal in the acceptor channel is observed. (C) Significant signal enhancement is observed in the FRET channel even before the data are processed to $FRET^{C}$ (from 18000 to 40000 a.u.).

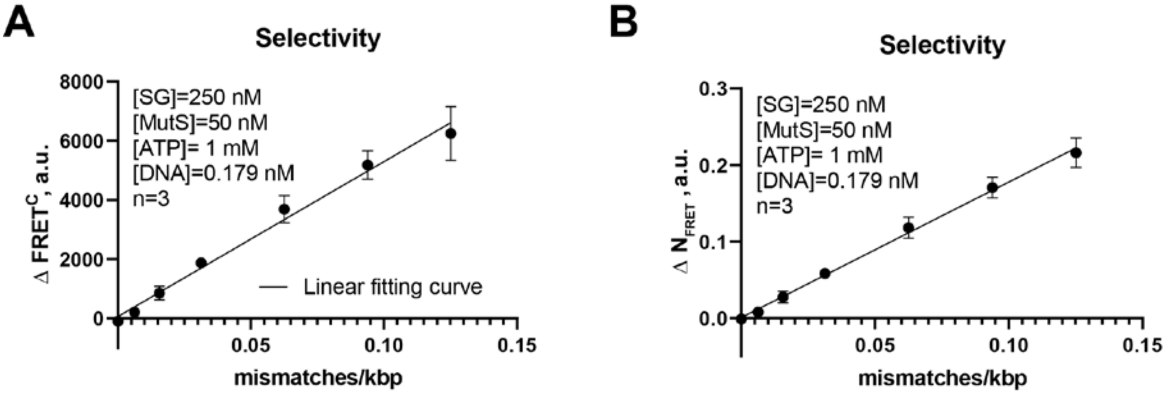


**Supplementary Figure S4.** Normalisation of the FRET signal decrease pipetting errors. In the case of a normalized signal, the signal intensities of both the donor and the acceptor are taken into account, resulting in smaller error bars (B) which further increases the selectivity of the method (equation 8).

$\Delta FRET^c$ was calculated as described in Figure 2C for the difference in $FRET^c$ before and after addition of ATP. Similar $\Delta N_{FRET}$ was calculated from the difference in $N_{FRET'}$ (calculated a described in equation 8.

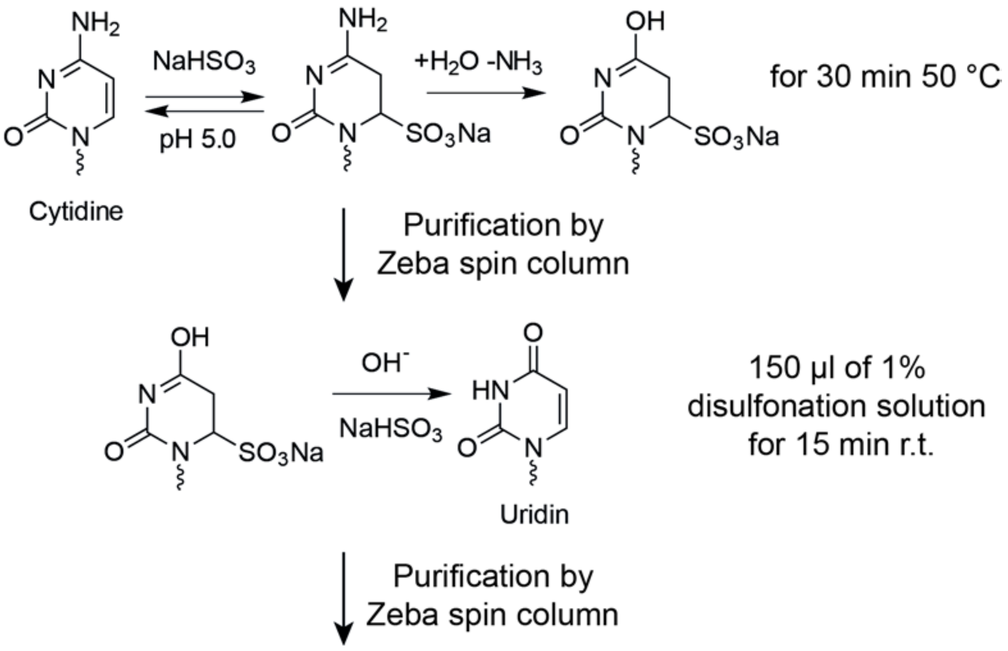


**Supplementary Figure S5. Deamination procedure**

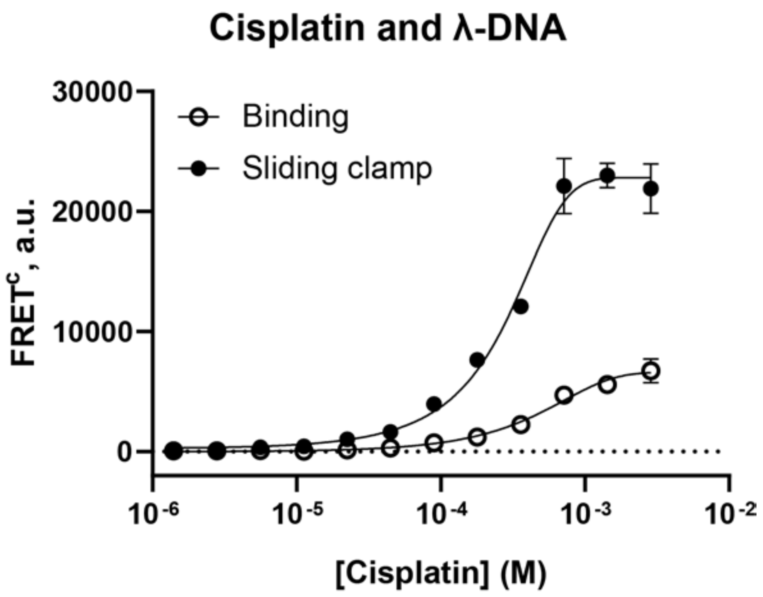


**Supplementary Figure S6.** Cisplatin treatment of the λ-DNA. Modification of λ-DNA and experiments with MutS were carried out as described in section “Cisplatin modification” for plasmid DNA.
